# Comprehensive population-based genome sequencing provides insight into hematopoietic regulatory mechanisms

**DOI:** 10.1101/067934

**Authors:** Michael Guo, Satish K. Nandakumar, Jacob C. Ulirsch, Seyedeh Maryam Zekavat, Jason D. Buenrostro, Pradeep Natarajan, Rany Salem, Roberto Chiarle, Mario Mitt, Mart Kals, Kalle Pärn, Krista Fischer, Lili Milani, Reedik Mägi, Priit Palta, Stacey B. Gabriel, Andres Metspalu, Eric S. Lander, Sekar Kathiresan, Joel N. Hirschhorn, Tõnu Esko, Vijay G. Sankaran

## Abstract

Genetic variants affecting hematopoiesis can influence commonly measured blood cell traits. To identify factors that affect hematopoiesis, we performed association studies for blood cell traits in the population-based Estonian Biobank using high coverage whole genome sequencing (WGS) in 2,284 samples and SNP genotyping in an additional ~17,000 samples. Our analyses identified 17 associations across 14 blood cell traits. Integration of WGS-based fine-mapping and complementary epigenomic data sets provided evidence for causal mechanisms at several loci, including at a novel basophil count-associated locus near the master hematopoietic transcription factor *CEBPA.* The fine-mapped variant at this basophil count association near *CEBPA* overlapped an enhancer active in common myeloid progenitors and influenced its activity. *In situ* perturbation of this enhancer by CRISPR/Cas9 mutagenesis in hematopoietic stem and progenitor cells demonstrated that it is necessary for and specifically regulates *CEBPA* expression during basophil differentiation. We additionally identified basophil count-associated variation at another more pleiotropic myeloid enhancer near *GATA2*, highlighting regulatory mechanisms for ordered expression of master hematopoietic regulators during lineage specification. Our study illustrates how population-based genetic studies can provide key insights into poorly understood cell differentiation processes of considerable physiologic relevance.

## INTRODUCTION

The human hematopoietic system is among the best understood paradigms of cell differentiation in physiology (1). Yet, despite our sophisticated understanding, many aspects of this process remain poorly understood. In particular, while hematopoiesis is perturbed in a variety of human blood disorders and shows considerable inter-individual variation, the underlying basis of the disease etiology and variation remains incompletely understood. Genetic variation in hematopoiesis can be reflected in commonly measured laboratory values, such as hemoglobin levels or blood cell counts. Rare mutations disrupting genes involved in hematopoiesis can result in severe anemias or other blood disorders, which can present with significant reductions in blood cell counts (2). Common genetic variants affecting hematopoiesis can also subtly influence blood cell measurements in the general population and can alter the clinical manifestations in rare blood disorders (1, 3, 4). Genetic studies offer a unique opportunity to gain insight into the hematopoietic system without being biased by our prior knowledge.

The Estonian Biobank is a population-based biobank that has collected DNA samples from 51,535 individuals representing approximately 5% of the Estonian population (5). This cohort is composed of adults representative of the larger Estonian population in terms of age, sex, and geographic distribution. The biobank is unique in that electronic medical records (EMRs) in Estonia are centralized and all participants have consented to allow full access to their medical records, which provides an excellent resource to investigate the underlying genetic basis for a variety of traits and diseases. Moreover, many of the samples from the biobank have undergone extensive genomic characterization, including single nucleotide polymorphism (SNP) genotyping from ~17,000 individuals and PCR-free, high-coverage whole genome sequencing (WGS) from 2,284 individuals. Here, in order to gain insight into hematopoiesis and regulatory mechanisms underlying this process, we have taken advantage of the valuable resource afforded by the Estonian Biobank to perform genetic association studies of all blood cell measurements available in this large population-based cohort.

## RESULTS

### Study overview

To perform the genetic association studies for blood cell traits, we used the WGS of 2,284 individuals and the SNP genotypes of ~17,000 individuals from the Estonian Biobank. The WGS data underwent joint variant calling, followed by extensive sample and variant-level quality control (QC) (Dataset S1). The SNP genotypes were used to impute genotypes to a custom reference panel constructed from the high coverage Estonian Biobank WGS data.

Using the genotype data described above, we tested for associations with 14 blood cell measurements. This included measurements reflective of red blood cell (RBC) numbers, size, and other related parameters (hemoglobin, hematocrit, RBC count, mean corpuscular volume (MCV), mean corpuscular hemoglobin (MCH), and mean corpuscular hemoglobin concentration (MCHC)); platelet numbers and size (mean platelet volume (MPV)); as well as white blood cell subtype numbers (absolute numbers of neutrophils, monocytes, lymphocytes, eosinophils, and basophils). For a subset of randomly selected individuals, blood cell measurements were directly assayed in a clinical laboratory (hereafter referred to as “lab-based”) (Dataset S2, Dataset S3 and Fig. S1). For the remaining individuals, we mined the EMR to extract blood cell measurements when available. Since all individuals in the Estonian Biobank consented to provide access to their corresponding EMR data, we were able to greatly expand sample sizes in a resource-efficient manner. We performed single variant association analyses on all variants with a minor allele count of greater than or equal to 3. We also performed gene-based burden testing of rare variants (MAF<5%) using SKAT-O (6).

### Blood cell trait associations in the Estonian Biobank

The single variant analysis revealed a total of 17 genome-wide significant associations (p<5x10^−8^) across the various blood cell measurements (Table 1). Sixteen of these associations had been identified previously and highlight important biological mechanisms, such as associations at the *HBS1L-MYB* locus that contains several variants showing pleiotropy with multiple blood cell measurements (Dataset S4) (3, 7, 8). This locus is of considerable interest, since the blood trait-associated variants within this region are associated with the severity of the major hemoglobin disorders, sickle cell disease and β-thalassemia (8–10). Other loci that we identified here contain well-known hematopoietic regulators such as *JAK2* (associated with platelet counts) (11, 12) and *F2RL2* (associated with MPV) (13). In contrast to the GWAS studies involving common variants, the gene-based burden testing (which seeks to aggregate rare variants in each gene) did not identify any significant associations (at p-value < 8.33 x10^−7^). Our sample is likely underpowered for rare-variant analysis, which is expected to require considerably larger sample sizes in the range of tens of thousands of individuals (14).

**Table 1.** Detailed summary of significant associations. Both the WGS and combined (WGS+SNP genotyping) p-values are listed. Effect sizes (per minor allele) are based on untransformed trait values in the WGS only. Variance explained is based on inverse normal transformed trait values in the WGS only. CS column shows the number of variants in the CS. CS+NDR column shows the number of CS variants overlapping an ATAC-seq NDR.

| Locus | Position | Ref/Alt | rsID | MAF | WGS P-value | Combined P-value | Effect Size | Variance Explained | Trait | Gene | CS | CS+NDR |
| --- | --- | --- | --- | --- | --- | --- | --- | --- | --- | --- | --- | --- |
| 19q13 | 33754548 | C/T | rs78744187 | 0.104 | $1.25 \times 10^{-14}$ | $6.19 \times 10^{-38}$ | -0.0059 (1000/uL) | 0.044 | Basophil Count | *CEBPA* * | 1 | 1 |
| 12q24 | 122216910 | A/G | rs11553699 | 0.100 | 0.0016 | $7.04 \times 10^{-20}$ | 0.048 (fL) | 0.011 | MPV | *WDR66* | 1 | 1 |
| 3p14 | 56849749 | T/C | rs1354034 | 0.317 | 0.0012 | $1.29 \times 10^{-14}$ | -0.033 (fL) | 0.012 | MPV | *ARHGEF3* | 1 | 1 |
| 10q21 | 65063844 | T/A | rs61855497 | 0.369 | 0.0065 | $3.72 \times 10^{-14}$ | -0.023 (fL) | 0.0085 | MPV | *JMJD1C* | 18 | 4 |
| 6q23 | 135423209 | T/C | rs9373124 | 0.308 | 0.00014 | $6.86 \times 10^{-14}$ | 0.073 (pg) | 0.010 | MCH | *HBS1L/MYB* | 19 | 7 |
| 6q23 | 135419631 | A/G | rs9389268 | 0.304 | 0.0014 | $1.23 \times 10^{-13}$ | 0.19 (fL) | 0.0070 | MCV | *HBS1L/MYB* | 21 | 8 |
| 6q23 | 135419636 | C/T | rs9376091 | 0.304 | 0.011 | $6.96 \times 10^{-12}$ | -0.021 ($10^6$/uL) | 0.0044 | Red Blood Cell Count | *HBS1L/MYB* | 20 | 10 |
| 3q21 | 128296273 | G/A | rs2465283 | 0.101 | 0.0014 | $9.99 \times 10^{-12}$ | -0.0028 (1000/uL) | 0.0077 | Basophil Count | *GATA2* | 11 | 1 |
| 7q22 | 106370644 | C/G | rs342292 | 0.487 | 0.00062 | $3.26 \times 10^{-11}$ | 0.032 (fL) | 0.013 | MPV | *PIK3CG* | 46 | 6 |
| 9q31 | 113918856 | A/G | rs10980802 | 0.491 | $3.04 \times 10^{-6}$ | $5.06 \times 10^{-11}$ | -0.020 (1000/uL) | 0.016 | Monocyte Count | *LPAR1*<sup>c</sup> | 18 | 0 |
| 9p24 | 4763491 | G/A | rs12005199 | 0.332 | 0.028 | $1.17 \times 10^{-9}$ | 2.8 (1000/uL) | 0.0033 | Platelet Count | *JAK2* | 1 | 1 |
| 3p14 | 56849749 | T/C | rs1354034 | 0.319 | 0.015 | $4.40 \times 10^{-9}$ | 2.6 (1000/uL) | 0.0040 | Platelet Count | *ARHGEF3* | 1 | 1 |
| 6q23 | 135431640 | T/C | rs9494142 | 0.254 | $2.16 \times 10^{-5}$ | $7.51 \times 10^{-9}$ | 5.3 (1000/uL) | 0.012 | Platelet Count | *HBS1L/MYB* | 22 | 7 |
| 11p15 | 242859 | A/G | rs55781332 | 0.264 | 0.043 | $1.42 \times 10^{-8}$ | -0.022 (fL) | 0.0047 | MPV | *PSMD13* | 35 | 6 |
| 6p21 | 33545125 | A/G | rs5745587 | 0.274 | 0.0011 | $2.81 \times 10^{-8}$ | 4.1 (1000/uL) | 0.0072 | Platelet Count | *BAK1* | 23 | 2 |
| 5q13 | 75935631 | A/T | rs114685606 | 0.0230 | 0.53 | $3.41 \times 10^{-8}$ | 0.011 (fL) | 0.00046 | MPV | *F2RL2* | 3 | 1 |
| 22q12 | 37470224 | T/C | rs2413450 | 0.459 | 0.0052 | $3.58 \times 10^{-8}$ | 0.060 (pg) | 0.0054 | MCH | *TMPRSS6*<sup>c</sup> | 5 | 0 |

\*indicates novel locus. <sup>c</sup>Indicates presence of a coding variant in CS

The strongest effect identified was a novel association with basophil counts near *CEBPA* (rs78744187, p-value = 6.19x10^−38^) (Fig. 1 and Fig. S2). Each minor allele of this SNP is associated with a 5.9 (per μL) decrease in basophil counts and the SNP remarkably explains 4.4% of phenotypic variance (Table 1). To ensure that this association is not driven by extreme or spurious values from hospital-based measurements, we validated the association using only lab-based measurements with outliers removed (p = 3.31x10^−15^). Furthermore, to ensure that this association is not population-specific, we examined this SNP in 7,488 individuals from 3 US-based European ancestry cohorts and observed a significant association with basophil counts (p-value = 5.99x10^−7^; Fig. S3A). Despite the remarkably large effect size of this SNP, previous GWAS for basophil counts have not detected this association (15–17). This is likely because previous studies were imputed to a sparser reference panel (HapMap). Since none of the variants present in HapMap tag rs78744187 strongly, these studies would have failed to detect this association (Fig. S3B). This observation demonstrates how denser reference panels or comprehensive genome sequencing data can enable the discovery of additional common variants associated with human traits and diseases.

**Fig. 1.**
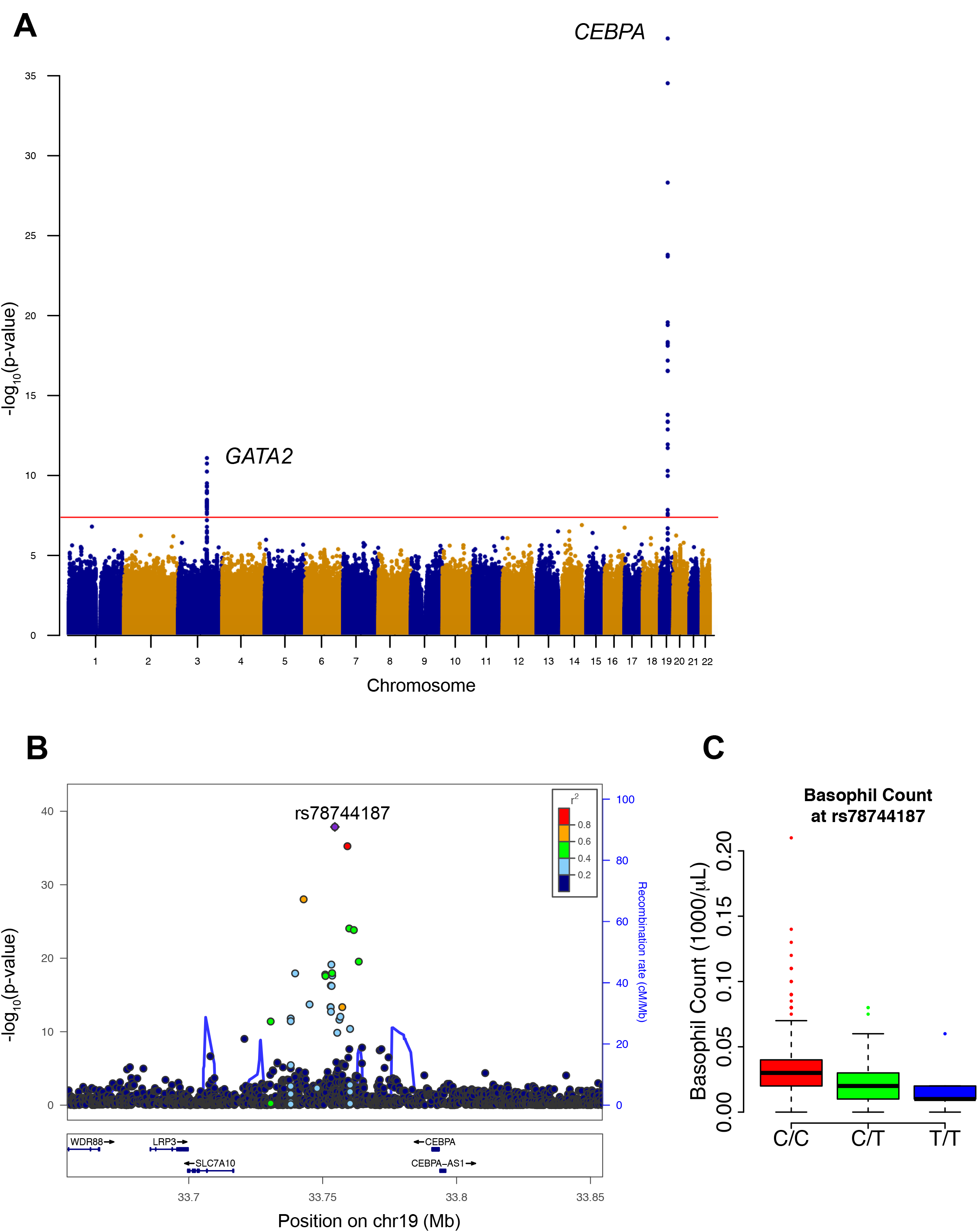
Novel basophil count association near *CEBPA.* (A) Manhattan plot for single variant association study for basophil counts. Genome-wide significant associations near *GATA2* and *CEBPA* are marked. (B) Locuszoom plot shows association strength, linkage disequilibrium, and recombination event frequency. (C) Basophil counts by genotype of rs78744187.

To assess the comprehensiveness of our analysis (Fig. S4), we compared all of the variants identified by genome sequencing at each locus with the 1000 Genomes (1000G) reference panel (18). While our study identified variants in significant linkage disequilibrium (LD) with the lead SNP (r^2^>0.5) that were absent from the 1000G phase 1 reference panel, all of these variants were present in phase 3 (19). Importantly, no variants identified in significant LD with the lead SNP (r^2^>0.5) in 1000G were missing from our analysis. Given these results, we were confident that all potentially causal variants had been captured by our analyses and our custom WGS-based reference panel was genuinely reflective of the study population, which are both important prerequisites for fine-mapping.

### Fine-mapping genetic associations

Although most of the associations have been previously detected, none have yet been pinpointed to specific variants. To attempt to identify the likely causal variant at each locus, we performed statistical fine-mapping analyses, which utilize LD patterns and association statistics to generate the probability that any particular variant at a locus of interest is causal. We applied three methods for fine-mapping (Approximate Bayes Factor (ABF), CaviarBF, and PICS) (20–22) and for each, generated a credible set (CS) of variants, which has a 97.5% probability of containing the casual variant. The CSs generated with ABF and CaviarBF exhibited near perfect concordance, whereas the CSs generated with PICS, while in strong agreement at most loci, included substantially more variants for 3 of the loci (Fig. S5). As these additional variants nominated solely by PICS were generally of low r^2^ to the sentinel association, the intersection of ABF and CaviarBF was chosen as the final CS. Remarkably, at four of the thirteen independent loci (MPV/ platelet counts at 3p14, platelet counts at 9p24, MPV at 12q24, and basophil counts at 19q13), our fine-mapping results resolved the association signal to a single putative causal variant (PCV). At two other loci, the CSs had three and five variants (Table 1 and Dataset S5). Thus, by resolving association signals to a finer resolution, we are able to generate experimentally tractable hypotheses about potential causal mechanisms, as we discuss in detail below. For the remaining ten associations, the CSs have between 11 and 46 variants (median of 20).

### Overlap with epigenomic data suggests causal mechanisms

An estimated 80-90% of causal GWAS signals are non-coding variants that presumably act by altering expression of nearby genes (23, 24). To define potential causal mechanisms of the variants, we overlapped CS variants with nucleosome depleted regions (NDRs) identified by ATAC-seq from 13 primary human cell types (25), comprising the majority of the hematopoietic hierarchy. For 35 CS variants (out of 186 variants from 17 loci; 18.8%), we identified an overlap with hematopoietic NDRs, a significant enrichment as compared to non-CS variants in moderate to high LD (r^2^ > 0.5) (OR = 2.41, p-value = 0.0009). Additionally, a permutation test involving local shifting of the NDRs around the CS variants revealed a significant enrichment (1.87-fold change in overlap, p-value = 0.00042) (26). Furthermore, at 11 out of 13 independent loci (85%), at least one CS variant overlapped a NDR (Fig. 2A). Of note, only the remaining 2 loci contained a coding variant in their CSs (Table 1).

**Fig. 2.**
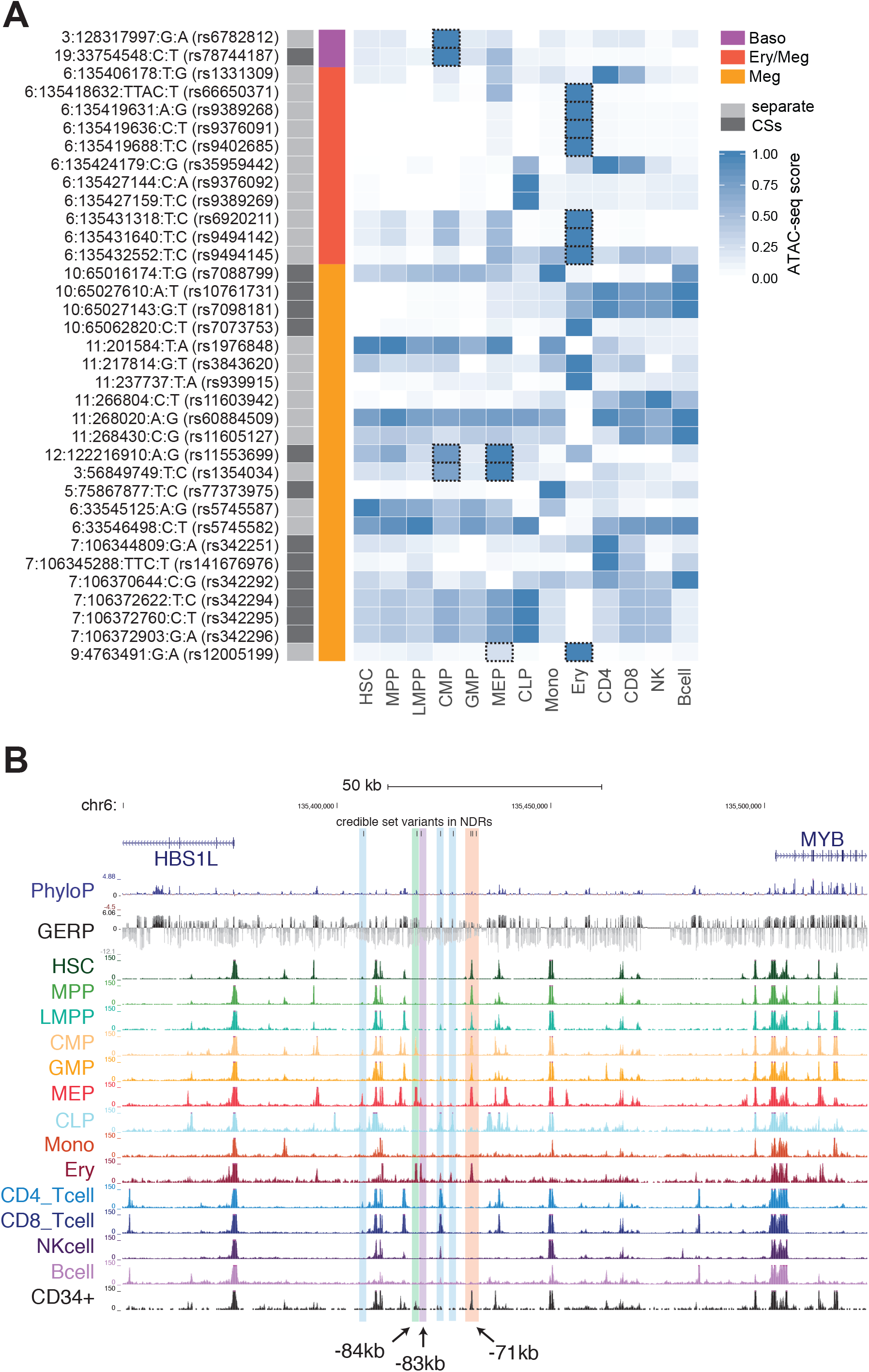
Integration of ATAC-seq data with fine mapping results sheds mechanistic insights. (A) Credible set variants that overlap with NDRs in the 13 hematopoietic cell types are shown. Quantile-normalized read counts per million were min-max scaled for each row. Based upon manual investigation, variants that overlapped with a NDR that was not within the top 20% of NDRs for at least one of the 13 cell types were excluded. Variants that fall within lineage-specific NDRs of clear relevance to their associated phenotypes are highlighted with dashed boxes. (B) 11 of the variants in the combined credible set for the *HSB1L/MYB* locus association with multiple red cell and platelet associations lie within 6 separate hematopoietic enhancer elements. The three MEP-/ erythroid-specific elements are shown in green (−84kb), purple (-83kb), and red (−71kb). Rs9494145 resides within a weaker −70kb element and is included in the same highlight as the substantially more nucleosome free −71kb element.

For example, at the well-known *HBS1L-MYB* locus (7), 11 variants associated with multiple red blood cell and platelet traits overlap with a NDR in at least one stage of hematopoiesis. Seven of these variants overlap with predominately erythroid-specific NDRs (Fig. 2A-B). While our results agreed with previous studies that variants in the −84kb and −71kb elements are putative functional variants (27), we also identified a previously uncharacterized −83kb erythroid element harboring 3 CS variants that may also have regulatory function (Fig. S6). Notably, for all four of the association signals that we fine mapped to a single variant, the identified variant overlaps with a hematopoietic NDR (Table 1). As an example, we were able to fine-map the association with MPV and platelet counts on 3p14 to rs1354034 which overlaps with a common-myeloid progenitor (CMP)- and megakaryocyte-erythroid progenitor (MEP)-specific regulatory element that may affect the transcription of *ARHGEF3*, a factor that has previously been implicated in hematopoiesis (Fig. 2A and Fig. S7) (28). In all of these examples, our comprehensive ascertainment of genetic variation gave us confidence that the causal variant is included among the variants we analyzed.

To further explore the putative regulatory modalities of these CS variants, we investigated the overlap of CSs with transcription factor (TF) occupancy, functional regulatory models, and predicted motif disruptions. Based upon functional models trained on TF occupancy, open chromatin, and histone modifications, CS variants were enriched for functional regulatory variants (Fig. S8) (29). Since TF occupancy profiles were not available for the entire hematopoietic hierarchy, we inferred putative TF overlap by investigating the overlap of CS variants with 4,559 publically available ChIP-seq datasets from human blood-based tissues and cell lines. These analyses revealed putative mechanisms for a number of variants and provide testable hypotheses, which are particularly tractable for the four CSs containing only a single variant (Dataset S6 and Dataset S7). For example, rs1354034, which we described above as being within a NDR near *ARHGEF3*, disrupts a conserved GATA motif. In addition, Gata1 occupies the orthologous mouse region containing this variant in megakaryocytes, but not erythroid cells, suggesting a putative mechanism by which this variant may act (Fig. S9).

### Basophil associations illuminate mechanisms for hematopoietic lineage specification

We next turned to the novel association with basophil counts at 19q13 near *CEBPA.* As we noted above, this locus could be resolved to a single PCV, rs78744187, which resides 39 kb downstream from *CEBPA*, near a separate +42kb enhancer that has been shown to influence *CEBPA* expression along various myeloid lineages (30–32). Rs78744187 appeared to be solely associated with basophil counts and showed no evidence of pleiotropic effects on other blood cell traits, including among other myeloid lineages (Dataset S4). Conditioning on rs78744187 suggested only one independent signal at this locus (Fig. S10A). Interestingly, rs78744187 resides within a distinct NDR present only in CMPs, but not in granulocyte-monocyte progenitors (GMPs), consistent with emerging data for a GMP-independent origin for basophils, mast cells, eosinophils, and their progenitors (Fig. 3A) (33–36). Moreover, this NDR is weakly to moderately present in myeloid cell lines from mice and humans (HL60, K562, HPC7, and CMK) and is occupied by numerous myeloid transcription factors, including master regulators of myeloid differentiation: GATA2 and RUNX1 (Dataset S6 and Fig. S11). In a luciferase reporter assay, the +39kb region demonstrated enhancer activity (~40 fold increase in activity relative to the minimal promoter) in the K562 myeloid cell line. Additionally, the basophil count-decreasing rs78744187-T allele was associated with a 1.4-fold reduction in enhancer activity (Fig. 4A). Despite extensive analyses, we were unable to elucidate an exact mechanism by which this variant affects transcription using our TF occupancy and predicted motif analyses, as is frequently noted to be the case for PCVs that alter gene expression (4, 22). Taken together, these data show that the +39kb region contains a myeloid enhancer element that is active in CMPs and that shows variation in activity modulated by the rs78744187 variant.

**Fig. 3.**
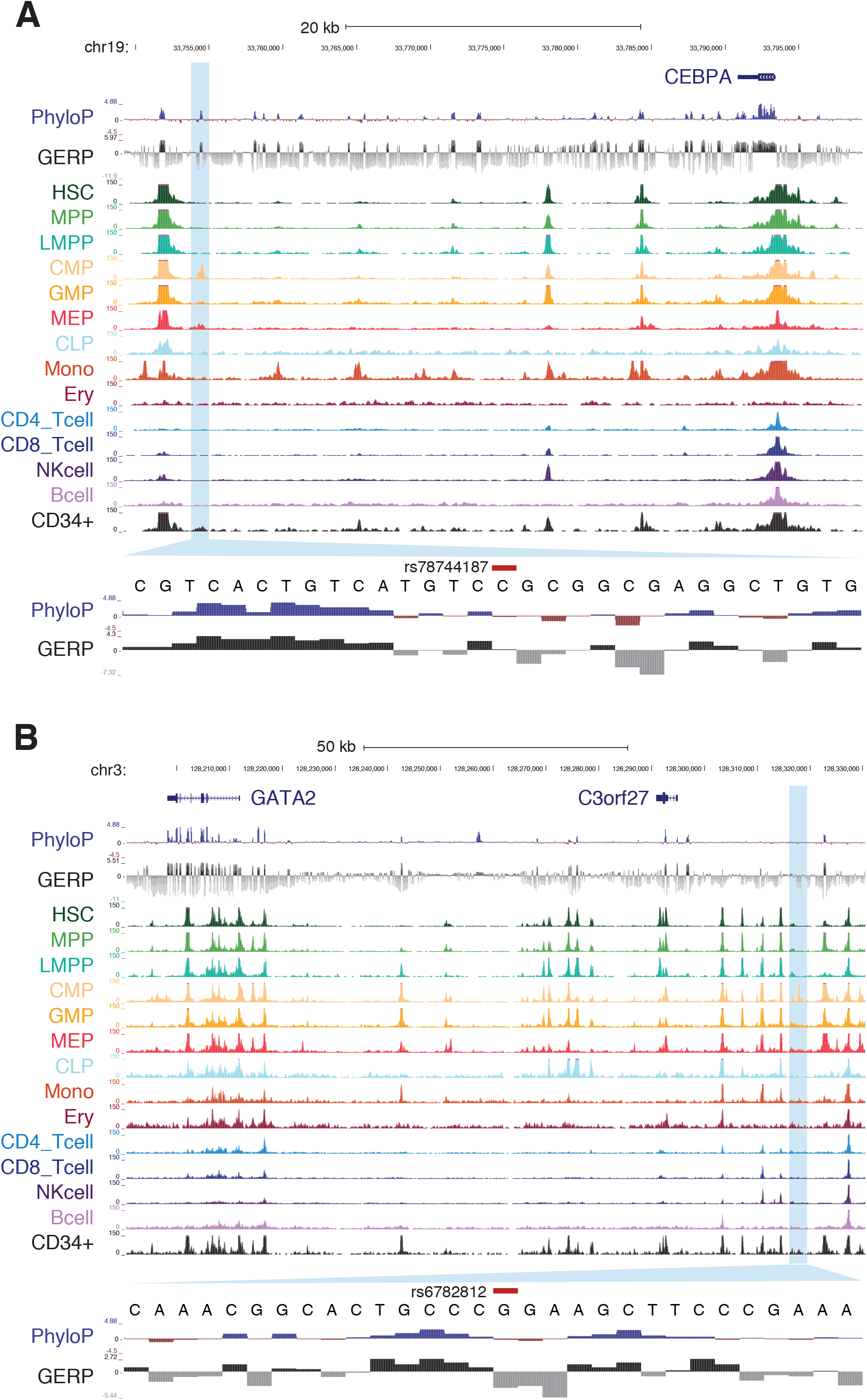
Overlap of basophil-associated variants with hematopoietic regulatory elements. (A) Overlap of rs78744187 with NDRs in hematopoietic progenitors and their terminal progeny. Conservation across 100 vertebrates (PhyloP) or mammals (GERP) is also shown. A conserved motif element is observed proximal to rs78744187. (B) Similar to (A) except for rs6782812. Two conserved motif elements can be observed nearby.

**Fig. 4.**
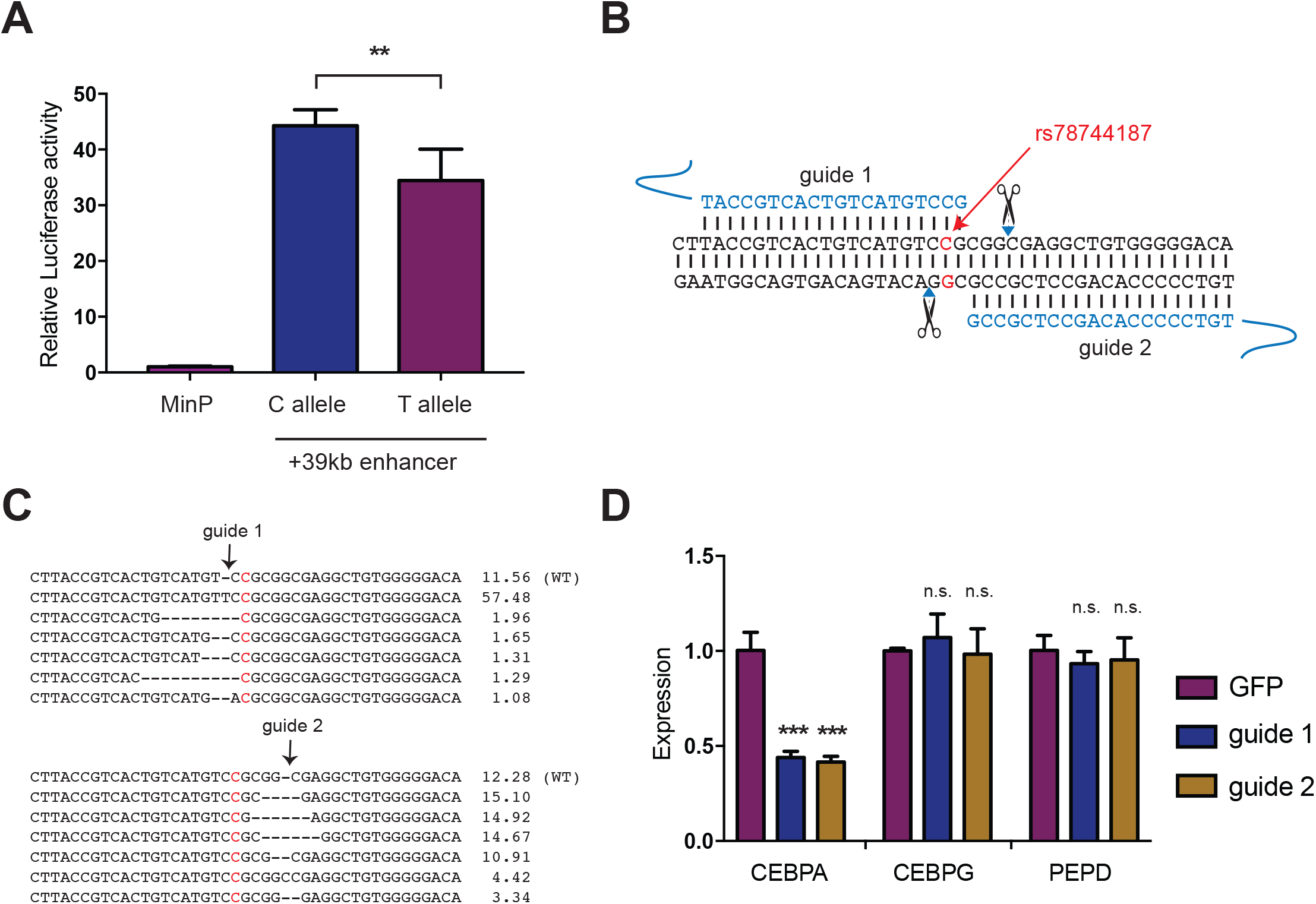
rs78744187 modulates the activity of a *CEBPA* enhancer. (A) A 400bp genomic region containing rs78744187 shows allele specific enhancer activity in K562 cells by luciferase assay (**, p-value < 0.01). (B) Schematic of CRISPR/Cas9 disruption at the +39kb myeloid enhancer disruption. (C) Mobilized peripheral blood CD34+ cells were infected with lentiviral CRISPR/Cas9 constructs. Indel frequency was measured at day 14 by deep sequencing and the top 6 indels are shown. (D) Expression of transcribed genes in the TAD containing rs78744187 after enhancer disruption at day 7 (qRT-PCR). Results are reported as mean and SD across three independent experiments (n.s., not significant; ***, p-value < 0.0001).

To identify the gene(s) whose expression is modulated by rs78744187 to influence basophil production, we performed *in situ* perturbation of the +39kb enhancer using CRISPR/Cas9-mediated mutagenesis in CD34^+^ human hematopoietic stem and progenitor cells (HSPCs). We targeted the +39kb enhancer using two guides that flank the rs78744187 variant (Fig. 4B). Deep sequencing of the target regions showed a high cutting efficiency (~88%) for both guides (Fig. 4C). We observed a 60% reduction in *CEBPA* expression in the non-clonal population of enhancer-disrupted hematopoietic cells compared to controls (Fig. 4D). As enhancer-promoter looping interactions primarily occur within topologically associated domains (TADs), we investigated all other expressed genes in the TAD harboring this variant but did not observed any significant changes in their expression (Fig. 4D and Fig. S12) (30). Interestingly, perturbation of this element in the granulocyte/monocyte cell lines HL60 and U937 did not result in any major alteration of *CEBPA* expression (Fig. S13), demonstrating the specificity of this regulatory element during basophil differentiation.

The TF CEBPA has been previously implicated in basophil specification, in particular during the bifurcation from the developmentally-related mast cell lineage (37–39). However, CEBPA is also implicated more broadly in hematopoiesis as a master TF (30, 40, 41), suggesting that its expression is temporally regulated to specify basophils and other terminal lineages (42). To test whether the +39kb enhancer provides this temporal regulation of *CEBPA* expression for proper basophil differentiation, we performed directed differentiation of the CRISPR/Cas9 enhancer-mutagenized HSPCs in the presence of IL-3 (43–45). The enhancer-mutagenized cells showed a significant reduction in basophil production based upon cell surface markers and morphology, as well as a proportionate increase in mast cells compared to controls (Fig. 5A-D and Fig. S14). In addition, the basophils produced in the enhancer-mutagenized cells frequently showed impaired maturation with a paucity of basophilic granules and a high frequency of empty or eosinophilic granules instead (Fig. 5C-D). These results demonstrate that an intact +39kb enhancer is required for proper expression of *CEBPA* during basophil differentiation and maturation. Our results also extend earlier studies in mice that suggested a key role for *Cebpa* in modulating the basophil/ mast cell lineage fate choice (46, 47).

**Fig. 5.**
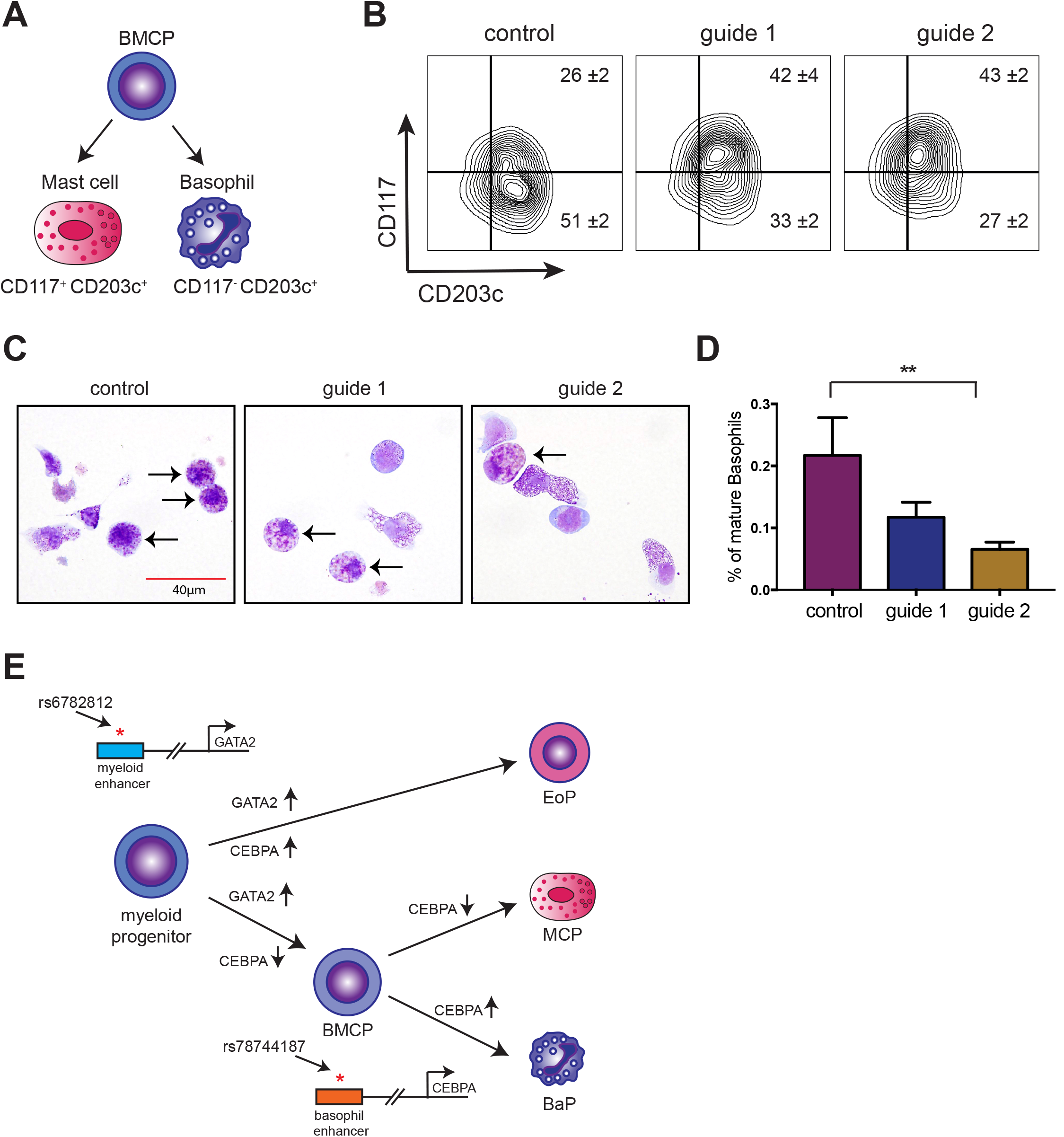
Intact +39kb CEBPA enhancer is required for human basophil differentiation. (A) IL-3 mediated differentiation of primary human CD34+ cells generates both basophils and mast cells from a common basophil/ mast cell progenitor (BMCP). (B) FACS analysis shows impaired differentiation of basophils and a concomitant increase in mast cells after +39kb enhancer disruption (mean +/− SD of three independent experiments). (C) Representative images of May-Giemsa stains at day 14. Arrows indicate fully differentiated, mature basophils in the left panel, while the arrows indicate cells with abnormal basophilic and some eosinophilic granules in the +39kb enhancer-disrupted cultures. (D) Impaired maturation of basophils based upon morphology in May-Grunwald Giemsa stains. Student’s t-test performed between control vs. both guides (**, p-value < 0.01). (E) Previous studies have shown that ordered expression of GATA2 and CEBPA is critical for differentiation of eosinophils, basophils and mast cells. Our GWAS follow-up study has identified enhancers that, at least partially, mediate this ordered expression pattern. Upregulation of GATA2 is required for all three lineages. Accordingly, the rs6782812 variant in the *GATA2* locus is associated with both eosinophil and basophil counts. Upregulation of CEBPA is required only for basophil differentiation from BMCPs and/ or CMPs. Accordingly, the rs78744187 variant in the *CEBPA* locus is associated with basophil counts and affects basophil differentiation. BMCP – basophil/mast cell progenitor, EoP – eosinophil progenitor, BaP – basophil progenitor and MCP – mast cell progenitor.

Our GWAS also identified an association with basophil counts at 3q21, which includes another master TF *GATA2* (rs2465283, p-value = 9.99x10^−12^) (Fig. 1A and Table 1). A previous GWAS performed in a Japanese population also identified an association at this locus with basophil counts (rs4328821; p-value = 5.3x10^−40^) (16). We noted that the basophil-decreasing allele of rs4328821 is associated with decreased *GATA2* expression in whole blood (p-value = 5.3x10^−13^) (48). By leveraging differences in the LD patterns between Estonians and East Asians and examining only variants in strong LD (r^2^>0.8) with the lead SNP in both populations, we were able to reduce our CS from 11 to 6 variants. Of these 6 CS variants, only one variant (rs6782812) overlapped a strong hematopoietic NDR. Surprisingly, similar to rs78744187, this NDR is also CMP-specific (Fig. 3B) and is occupied by the RUNX1 and GATA2 TFs (Fig. S15). In luciferase-based assays in K562 cells, the NDR demonstrates ~4.5 fold enhancer activity and rs6782812 reduced enhancer activity by 69% (Fig. S15A). Because there are common master TFs at both variant-harboring enhancers at rs2465283 *(GATA2)* and rs7874418 *(CEBPA)*, we examined whether these two variants might show an epistatic interaction. We found no evidence of epistasis between rs2465283 and rs7874418 (p-value = 0.070). The *GATA2-* associated variant was also associated with eosinophil counts (p-value = 3.07x10^−3^; Dataset S4), as has been seen previously in other studies (15–17, 49). An independent association near *GATA2* for monocyte counts has been reported by other studies (monocyte sentinel SNP rs9880192; r^2^=0.054 to rs2465283 in Europeans) (15, 17). These associations near *GATA2* are consistent with the well-known role of GATA2 in driving myeloid differentiation (50, 51). Together, these results suggest that rs6782812 influences lineage specification at an earlier myeloid progenitor that is capable of producing basophils, eosinophils, and potentially other lineages, while the *CEBPA* variant appears to be present in an enhancer that is specifically necessary for production of basophils from a downstream bi-potential basophil/ mast cell progenitor (BMCP) (Fig. 5E).

### Examination of disease associations

Basophils have been implicated in inflammation and host defense, but the causal role that basophils play in human disease is poorly understood (37, 52–54). To identify potential disease roles for basophils, we performed a phenome-wide association study (pheWAS) for rs78744187 and rs2465283 (55). We tested for the existence of associations between either variant and 534 diseases that had greater than 100 cases using associated ICD10 medical billing codes. No disease associations reaching the p-value threshold of 9.2x10^−5^ (following Bonferroni correction) were identified for either SNP. However, rs78744187 was nominally associated with joint derangements and enteropathic arthropathy (p-values of 0.00023 and 0.00059, respectively), which may have autoinflammatory etiologies (Dataset S8). Previous studies have identified multiple associations with inflammatory bowel disease (IBD) near *CEBPA* (56–58). However, the basophil association at rs78744187 appears to be independent from the IBD associations (Dataset S9). Additionally, the *GATA2* signal has been previously associated with risk of asthma, which was reported to be due to its role in regulating eosinophil counts (49). We could not replicate this asthma association at *GATA2* in our study (p-value = 0. 33). We do note that while disease associations with basophil counts are likely to exist, similar to those seen with the related eosinophil lineage, we are likely to be underpowered in our current study to robustly detect such an effect, particularly given the variable fidelity of medical coding (59).

## DISCUSSION

In this study, we integrated WGS-based genome-wide association studies, fine-mapping, epigenomic datasets, and functional assays to provide additional insight into our evolving understanding of lineage specification during human hematopoiesis (60, 61). Integration of comprehensive genetic and extensive epigenomic data at these loci provided key insight into human hematopoietic regulatory mechanisms. For example, we were able to identify a variant that likely affects GATA1 TF binding to influence expression of *ARHGEF3* during megakaryopoiesis. At the extensively studied *HBS1L-MYB* locus, by overlapping fine-mapping data with extensive ATAC-seq data, we provided evidence for additional putative causal variants. By integrating these complementary datasets, we were able to generate experimentally tractable hypotheses for further functional investigation.

At one of these loci, we fine-mapped a novel association with basophil counts near the master TF *CEBPA* to a CMP-specific enhancer element. Functional assays revealed that the causal variant altered enhancer activity and resulted in decreased *CEBPA* expression, which therefore helps drive the lineage choice between basophils and mast cells. In the region of another master TF, *GATA2*, our study identified a basophil and eosinophil count-associated variant within a similar CMP-specific enhancer associated with *GATA2* expression. Thus, our study provides evidence that common genetic variation regulates basophil production by tuning the ordered expression of master TFs through the alteration of stage-specific enhancer elements (42). Furthermore, as both basophil-associated variants fall within enhancer elements that are active specifically in CMPs (but not GMPs or MEPs), our study provides strong support for revised models of hematopoiesis, where eosinophils, basophils, mast cells, and their progenitors bifurcate at the earlier CMP stage, rather than the more traditional models where these lineages arise from GMPs along with granulocyte and monocyte progenitors (Fig. 5E) (35, 36). Our findings provide key insights into the molecular regulation of basophil production, an important and non-redundant cell type in inflammation and host defense that has been challenging to study in humans due to its rarity (37, 52–54). The identification of these variants will also allow for further studies of the mechanisms by which genetic variants influencing basophil counts may impact on human diseases.

Our study also demonstrated the benefits of using high coverage WGS in a population-based biobank. Comprehensive ascertainment of genetic variation allowed us to identify the novel association near *CEBPA*, which would have been missed had we imputed to sparser reference panels, such as HapMap. Furthermore, the high coverage WGS allowed us to comprehensively capture variation that might be missed by lower coverage sequencing approaches (such as longer indels and variants in low complexity regions), giving us confidence that the true causal variant has been identified at each locus, an important prerequisite for fine-mapping. Moreover, by performing our study in a population-based biobank, we were also able to link genetic data with EMRs to greatly increase sample sizes in a resource-efficient manner, providing support for similar programs such as the Precision Medicine Initiative (62). Together, our study demonstrates how key genetic and biological insights can be gained from comprehensive genetic studies in population-based biobanks.

## MATERIALS AND METHODS

### Blood cell measurements

For a subset of 1018 randomly selected individuals, we performed a complete blood count (CBC) in a clinical laboratory (“lab-based”). Clinical lab-based measurements where performed at Tartu University Clinic’s Diagnostics center. More details on specific methods and equipment used can be found here (http://www.kliinikum.ee/yhendlabor/analyysid). For remaining individuals, we extracted blood cell measurements from hospital-based records as available. Hospital based cell measurements were obtained through linkage with two main hospitals in Estonia (Tartu University Clinic and Northern Estonia Regional Hospital) and systematically mining Electronic Health Records for blood cell measurements. Presence of other diseases was not taken into account when normalizing the blood cell measurements.

For each individual, we used the hospital-based measurements only if lab-based values were not available. We removed spurious values and extreme outliers (Dataset S2). We then performed regression using a linear mixed model adjusting for sex as a fixed effect, and setting (hospital-based versus laboratory, as well as the specific hospital/clinic) and age at measurement as random effects. We took the median residual for each individual and performed inverse normal transformation of the median residuals. These median residuals were used for downstream association analyses.

Generation of genome sequencing data, variant calling, imputation, and association testing are all described in SI Materials and Methods.

### Luciferase reporter assays

The genomic region containing major and minor allele of the variants rs78744187 (~400bp) and rs6782812 (~364bp) was synthesized as gblocks (IDT Technologies, Table S10) and cloned into the Firefly luciferase reporter constructs (pGL4.24) using BglII and XhoI sites. The Firefly constructs (500ng) were co-transfected with pRL-SV40 Renilla luciferase constructs (50ng) into 100,000 K562 cells using Lipofectamine LTX (Invitrogen) according to manufacturer’s protocols. Cells were harvested after 48 hours and the luciferase activity measured by Dual-Glo Luciferase Assay system (Promega).

### Genome editing in human CD34+ HSPCs using lentiviral CRISPR/Cas9 mutagenesis

Two guide RNAs targeting the variant rs78744187 and a control guide RNA targeting GFP (Fig. 4B) were cloned into LentiCRISPRv2 constructs^21^. The constructs along with packaging helper constructs were transfected into HEK-293T cells for lentiviral production. The viral supernatant was then concentrated 60X by ultracentrifugation. Human CD34+ hematopoietic stem/progenitor cells (adult) were purchased from Seattle Fred Hutchinson Center and cultured in IMDM with 10% Fetal Bovine Serum in the presence of human Interleukin-3 (10ng/ul). On day 2 in culture ~500,000 cells were spinfected with the concentrated lentiviral supernatant and polybrene (8μg/ml) on retronectin coated plates (Takara). On days 5 and 6 in culture the cells were selected with puromycin (1μg/ml). *CEBPA* expression was measured at day 7 in culture. The cells were subsequently cultured until day 14 for differentiation into basophils and mast cells.

### Genome editing in HL60s and U937 cell lines using lentiviral CRISPR/Cas9

HL60 and U937 cells were cultured in RPMI with 10% Fetal Bovine Serum. 1-2 million cells were spinfected with lentiviral supernatant with polybrene (8μg/ml). On days 3, 4 and 5 post-spinfectio,n cells were selected with puromycin (1μg/ml). *CEBPA* expression was measured at day 12 post-spinfection. For the Surveyor assay, genomic DNA was extracted at day 12 and a 600bp region containing the CRISPR cut sites was PCR amplified (Dataset S10). The Surveyor assay was performed according to kit recommendations (IDT Technologies).

### Flow cytometry

Cells were incubated with Human BD Fc Block (BD Biosciences) for 10 minutes at room temperature to prevent non-specific binding to Fc receptors. Subsequently the cells were stained with CD117-PE (Clone 104D2, Biolegend) and CD203c-APC (Clone NP4D6, Biolegend) antibodies and analyzed by BD Accuri Flow Cytometer. FACS plots were generated by FlowJo (TreeStar).

## ACKNOWLEDGEMENTS

We would like to thank members of the Sankaran and Hirschhorn Labs, as well as the Estonian Genome Center, and numerous colleagues for valuable comments and discussions. This work was supported by the National Institutes of Health grants R01 DK103794, R33 HL120791 (to V.G.S.), and R01 DK075787 (to J.N.H). P.P. was funded by the Nordic Information for Action eScience Center by NordForsk (Project number 62721). The Estonian Genome Center was supported by the Estonian Research Council (PerMed I, IUT20-60), European Union H2020 grants (692145, 676550, 654248), and European Union through the European Regional Development Fund (GENTRANSMED).

